# Presynaptic GABAB Signaling Enhances Synaptic Inhibition Without Mitigating Persistent Dentate Gyrus Hyperexcitability After Status Epilepticus

**DOI:** 10.64898/2026.09.20.752988

**Authors:** Deepak Subramanian, Shivakeshavan R. Giridharan, Aayma Irfan, Vijayalakshmi Santhakumar

## Abstract

The dentate gyrus (DG) is a critical regulator of cortical input processing in the hippocampus, supporting spatial navigation and memory processing. In epilepsy, the DG undergoes extensive structural and functional reorganization that diminishes its ability to filter afferent inputs, resulting in enhanced hippocampal hyperexcitability, seizures, and cognitive impairment. Yet, how DG network dynamics evolve over time in the epileptic brain is not fully understood. Here, we investigated DG network excitability, feedback inhibition, and throughput in rats 1 week, 8 weeks, and 4 months after pilocarpine-induced status epilepticus (SE), and examined the role of presynaptic GABA_B_ receptor-mediated modulation of DG activity. Recording from urethane-anesthetized rats *in vivo*, we show a persistent increase in excitation-spike (E-S) coupling after SE. In contrast, feedback inhibition measured by short-duration paired-pulse ratios (PPR at 20ms, 40 ms and 60ms intervals) was significantly increased 1 and 8 weeks after SE, but was indistinguishable from controls by 4 months. Notably, we found that dentate spike (DS) frequency remained persistently elevated up to 10 months post-SE. Selective inhibition of presynaptic GABA_B_ receptors in the DG using CGP36216 further enhanced E-S coupling at 1 and 8 weeks in post-SE rats without affecting age-matched controls. CGP36216 did not alter E-S coupling in rats 4 months after SE, revealing a homeostatic role for basal presynaptic GABA_B_ signaling in restraining network excitability early after SE. CGP36216 robustly increased PPR at 20ms across all groups without altering DS frequency, suggesting that GABAB receptors boost DG feedback inhibition. Mechanistically, CGP36216 reduced single stimulus-evoked inhibitory postsynaptic currents (eIPSCs) in dentate granule cells without altering excitatory transmission after SE. Together, these findings support a disinhibitory role for presynaptic GABA_B_ receptors, which bolsters synaptic inhibition to transiently counter post-SE hyperexcitability, and offers a targetable mechanism for therapeutic intervention during epileptogenesis.

## INTRODUCTION

Epilepsy is a chronic neurological disorder characterized by hyperexcitable and hypersynchronous neuronal activity. The hippocampus, a key structure within the limbic system, possesses an innate network architecture that renders it highly susceptible to generating synchronized epileptiform activity. The hippocampal dentate gyrus (DG) serves as a critical regulatory “gate,” filtering afferent signals from the entorhinal cortex (EC) and controlling the activation of downstream hippocampal subfields. This gating mechanism allows the DG to transform diffuse cortical inputs into sparse, patterned downstream activation essential for cognitive processes such as spatial memory and pattern separation (Borzello et al., 2023; Hainmueller and Bartos, 2020; Leutgeb et al., 2007; McNaughton and Morris, 1987). During immobility and non-REM sleep, DG exhibits large-amplitude, synchronized population events called *dentate spikes* (DS) that are proposed to regulate downstream hippocampal CA3/CA1 activity (Bragin et al., 1995; Farrell and Soltesz, 2025; Headley et al., 2017; Penttonen et al., 1997). Through this regulatory control, DS play an important role in memory consolidation and recall (Ewell, 2017; Farrell et al., 2024; Lensu et al., 2019; Nokia et al., 2017)

The DG network output is tightly regulated by robust GABAergic inhibition, maintained by a highly divergent population of local interneurons (Freund and Buzsáki, 1996; Houser, 2007; Krook-Magnuson et al., 2015). In epilepsy, the DG undergoes extensive inhibitory cell loss and subsequent reorganization that profoundly diminishes the dentate gate (Heinemann et al., 1992; Lothman et al., 1992; Sloviter, 1987). Consistent with this, DS frequency have been shown to increase sharply during the early weeks following brain injury and kainic acid-induced status epilepticus (SE) (Flynn et al., 2015; Subramanian et al., 2026). Crucially, whether changes in DG network excitability, feedback inhibitory regulation, and throughput are transient or continue to persist/evolve over time following SE remains unknown.

In chronic epilepsy, the loss of interneurons and dysfunction of the surviving inhibitory pathways severely impair GABA_A_-mediated control of hippocampal circuits (Bernard et al., 2000; Fritschy et al., 1999; Sloviter, 1994a, 1987). While the role of fast synaptic inhibition in maintaining the dentate gate has been extensively evaluated, the contribution of metabotropic GABA_B_ receptors in shaping the dentate gate and its changes during epileptogenesis is understudied (Avoli and Lévesque, 2022). GABA_B_ receptors play an essential role in modulating the strength of EC-to-DG perforant path synapses, thereby influencing spatial and pattern learning (Brucato et al., 1996; Heaney and Kinney, 2016; Mott et al., 1993; Mott and Lewis, 1991). GABA_B_ receptors are densely expressed throughout the hilus and are targeted to both pre- and postsynaptic domains, a localization regulated, in part, by the 1a and 1b splice variant-specific sushi domains of GABA_B_ that direct axonal versus dendritic trafficking (Kulik et al., 2003; Vigot et al., 2006). Activation of presynaptic GABA_B_ heteroreceptors on interneurons has been shown to inhibit GABA release onto dentate granule cells (DGCs) and parvalbumin (PV)-positive basket cells (Foster et al., 2013; Mott et al., 1993; Yu et al., 2016). Thus, GABA_B_ receptors are positioned to act as a dynamic and cell-type-specific regulator of DG inhibition (Booker et al., 2020; Dugladze et al., 2013; Loureiro et al., 2025; Mott et al., 1999; Pelkey et al., 2017). Consequently, presynaptic GABA_B_ signaling can influence inhibitory microcircuit function and overall circuit excitability. Despite the availability of selective pharmacology for presynaptic GABA_B_ receptors, the role of presynaptic GABA_B_ receptors in dentate circuit function is unclear.

An increase in GABA_B_ receptor mRNA and protein expression is observed both in human hippocampal sclerosis tissue and in rodent models (Billinton et al., 2001; Furtinger et al., 2003; Princivalle et al., 2002; Straessle et al., 2003). A cell-type specific upregulation of presynaptic GABA_B_ auto-receptors in rodent kindling models was shown to enhance GABA_B_-mediated suppression of synaptic release, leading to altered disinhibition and aberrant gamma oscillations (Dugladze et al., 2013; Straessle et al., 2003; Yu et al., 2016). Since activation of presynaptic GABA_B_ heteroreceptors on interneurons inhibits GABA release, upregulation of this mechanism could further compromise the dentate gate and augment network throughput and excitability. Whether presynaptic GABA_B_ receptor-mediated regulation of synaptic inhibition contributes to the progressive changes in hyperexcitability seen during epileptogenesis remains an important knowledge gap.

In this study, we show that DG excitation-spike coupling and dentate spike frequency are persistently elevated after SE, indicating chronic dysregulation of DG circuit function and throughput. In contrast, polysynaptic feedback inhibition was transiently enhanced at 8 weeks post-SE, and was comparable to controls by 4 months. Selective presynaptic GABA_B_ receptor antagonism further enhanced DG excitation-spike coupling, indicating that GABA_B_ receptor plasticity early after SE may have a compensatory function. GABA_B_ receptor antagonism reliably enhanced the DG paired-pulse ratios without altering DS frequency in both control and post-SE rats. Consistent with a role for GABA_B_ receptors in enhancing synaptic inhibition after SE, selective antagonism of presynaptic GABA_B_ receptors reduced evoked IPSCs after SE. Together, these findings uncover a novel circuit-level bolstering of inhibition by presynaptic GABA_B_ receptors in the DG, possibly by polysynaptic disinhibition of GABAergic interneurons, that is observed early after SE and offers a targetable mechanism for therapeutic intervention.

## METHODS

### Animals

All experiments were performed in accordance with IACUC protocols approved by the University of California at Riverside, CA, and in keeping with the ARRIVE guidelines.

### Pilocarpine induced Status Epilepticus

Male Wistar rats (p25-28) were treated with Pilocarpine hydrochloride (330mg/kg; i.p) 20 min after Scopolamine Methylnitrate (1mg/kg) to induce status epilepticus (SE). Only rats that exhibited three or more convulsive seizures within the first hour after induction were included in the study. Diazepam (10mg/kg) was administered to stop SE after 60min. Saline-injected animals receiving identical scopolamine and diazepam treatments acted as controls. Animals were examined at 1 week, 8 weeks, and 4-10 months after SE induction.

### Stereological Estimation of Hilar Neurons

To quantify neuronal loss, control and post-SE rats (n = 4 rats/group, 1-month post-induction) were deeply anesthetized with Euthasol (150 mg/kg, i.p.) and transcardially perfused with PBS followed by 4% PFA. Brains were post-fixed and coronally sectioned (50 µm) using a compressotome (VF-310-Z, Precisionary Instruments). Coronal brain sections were processed for Nissl staining (0.1% cresyl violet acetate and 0.25% glacial acetic acid). Unbiased stereological estimates of hilar neuron populations were obtained using the Optical Fractionator probe (Stereo Investigator; MBF Bioscience) on a Zeiss Axioscope-5 microscope by blinded investigators as described previously (Subramanian et al., 2026). Cell counts from three-five slices were averaged within each animal and used for statistical analysis.

### In vivo electrophysiology

Rats were anesthetized using urethane (1.5g/kg i.p.) and head-fixed onto a stereotaxic frame. Under aseptic conditions, rats were implanted with a single wire electrode 50 µm tungsten wire depth electrode (California Fine Wire company, Grover beach, CA), attached to a cannula (P1 technologies, Roanoke, GA). The electrode-cannula assembly was positioned in the granule cell layer (AP: 3-4 mm, ML: 2 mm, DV: 3-3.5 mm) guided by electrical stimulation of the angular bundle (ML:4-4.2 mm, DV: 1.8-2 mm; pulse width 150µs delivered at 0.1Hz). Two screw electrodes on the contralateral side served as reference and ground. Dental cement was used to secure the electrode in position. Acute recordings from urethane-anesthetized rats started 60 minutes after induction and the depth of anesthesia was confirmed by toe pinch every 30 minutes. At the end of the experiment, animals were euthanized with a lethal dose of Euthasol (150 mg/kg, i.p.). Signals were amplified 100 times (Pinnacle Systems, Lawrence, KS), sampled at 10kHz, digitized using PowerLab 16/35, and recorded using LabChart8 software (AD Instruments, Colorado Springs, CO). Analysis was conducted using LabChart modules. To assess network excitability, we derived current-response curves by electrically stimulating the angular bundle (150 µs @ 0.1Hz). Population spike amplitude and field EPSP slope were analyzed using modules in LabChart software. Feedback inhibition was evaluated by quantifying responses to pairs of stimulus pulses at 20 ms, 40 ms, and 60 ms interpulse intervals. To assess network excitability and dentate spikes (DS), DG local field potentials (LFPs) were recorded in 15-minutes blocks with a series of 5 paired stimuli (20ms interval @ 0.1Hz) delivered at the start of each block (See Figure 4B). In some experiments, CGP36216 (3 mM; 0.5μl), a GABA_B_ antagonist selective for presynaptic receptors, or saline was injected into the DG at a constant rate of 0.1 μl/min using a Hamilton syringe connected to a syringe pump (11 Elite Nanomite, Harvard Apparatus).

Dentate spikes were recorded either using an electrode-cannula assembly (as described above) or 16 Ch silicon probe (A1x16-5mm-100-703 CM16, Neuronexus, Ann Arbor, MI). Probe recordings were amplified 100 times using a Model 3600 amplifier (AM-Systems, Sequim, WA), digitized using PowerLab 16/35, and recorded using LabChart 8 software (AD Instruments, Colorado Springs, CO). Dentate spikes from 16-channel probe recordings were detected using the Toothy analysis pipeline (https://github.com/Farrell-Laboratory/Toothy). Briefly, large positive events were identified using a 4.5 standard deviation (S.D) threshold. Detected events were visually confirmed by a blinded investigator, and the total number of DSs recorded in 30 minutes was quantified. To assess drug effects on DS, single-channel recordings of DG LFP were analyzed using custom Python code that detected events that exceeded a 4.5 S.D. threshold, with a half-width less than 25 ms. The total number of DSs recorded in a 14-min period was used to evaluate drug effects. Dentate spikes were not subclassified as type 1 or type 2.

### Ex vivo electrophysiology

Control and post-SE rats were anesthetized under isoflurane and decapitated. The brain was extracted, and horizontal slices of the ventral hippocampus (350 µm) were prepared using Leica VT1200S Vibratome (Wetzlar, Germany) in ice-cold high-sucrose artificial Cerebro-Spinal Fluid (Sucrose-aCSF) that contained (in mM) 60 NaCl, 110 sucrose, 26 NaHCO_3_, 5 glucose, 7 MgSO_4_, 3 KCl, 1.25 NaH_2_PO_4_, and 0.5 CaCl_2_. Slices were bisected and incubated at 32 ± 1°C for a minimum of 1 hour in a submerged holding chamber containing recording aCSF and subsequently held at room temperature. The recording aCSF contained (in mM) 126 NaCl, 3 KCl, 2 CaCl_2_, 2 MgSO_4_, 1.25 NaH_2_PO_4_, 26 NaHCO_3_ and 10 D-glucose. All solutions were saturated with 95% O_2_ and 5% CO_2_ and maintained at a pH of 7.4 for 2-6 h.

Ventral hippocampal slices were transferred to a submerged recording chamber and perfused with oxygenated aCSF at 33 ± 1°C. Whole-cell voltage-clamp recordings from DG granule cells were obtained under IR-DIC visualization with a Nikon Eclipse FN-1 (Nikon Corporation, Shinagawa, Tokyo, Japan) or Olympus BX50 (Olympus Corporation, Shinjuku, Tokyo, Japan) microscope, using 40X water-immersion objectives. Recordings were obtained using MultiClamp 700B amplifiers (Molecular Devices, Sunnyvale, CA). Glass microelectrodes (3–5 MΩ), pulled with a Sutter P-1000 Glass puller (Sutter Instruments, Novato, CA), were used to obtain recordings. Excitatory/inhibitory currents were recorded using a cesium-based internal solution (in mM: Cesium methanesulfonate 140, NaCl 5, HEPES 10, EGTA 0.2, Mg2ATP 2, Na2GTP 0.2, Qx314 5, (pH 7.25, 270-285 mOsm). Evoked excitatory postsynaptic currents (eEPSC) were recorded at a holding potential of -70 mV, and evoked inhibitory postsynaptic potentials (eIPSC) were recorded from the same dentate granule cells (DGCs) at a holding potential of 0 mV. Biocytin (0.2%) was included in the internal solution for post-hoc cell identification (Gupta et al., 2012; Swietek et al., 2016; Yu et al., 2016). Data were digitized using DigiData 1440A and acquired at 10kHz using pClamp10. Data from cells in which access resistance changes over 20% during recordings and recordings with access resistance above 25 MΩ were excluded from analysis.

### Statistical Analysis

Data are expressed as mean ± SEM for parametric data or as median and interquartile range (IQR) for non-parametric data. Group comparisons of hilar neuron counts and dentate spike frequency were performed using unpaired Student’s t-tests. For electrophysiological experiments involving multiple time points, groups, or pharmacological interventions, data were analyzed using two-way repeated-measures (RM) ANOVA or two-way ANOVA, followed by post-hoc testing where appropriate. Significance was defined as *P*<0.05. All statistical analyses were conducted using GraphPad Prism 10.0.1 (Boston, MA).

## RESULTS

To establish the validity of our post-SE model, we first performed stereological estimations of DG hilar neurons 4-weeks after pilocarpine-induced SE (Figure 1A–B). Unbiased counting confirmed a significant reduction in total hilar neuron estimates in post-SE rats compared to age-matched saline-injected controls (Total hilar cell estimates: Control: 28,293 ± 2,927; post-SE: 19,608 ± 1,560; N=4 rats each; *P*=0.0397, unpaired t-test, t=2.619, df=6; Figure 1A–C). This hilar cell loss is consistent with classic models of post-SE dentate network reorganization and spontaneous epileptogenesis (Sloviter, 1994b).

**Figure 1:**
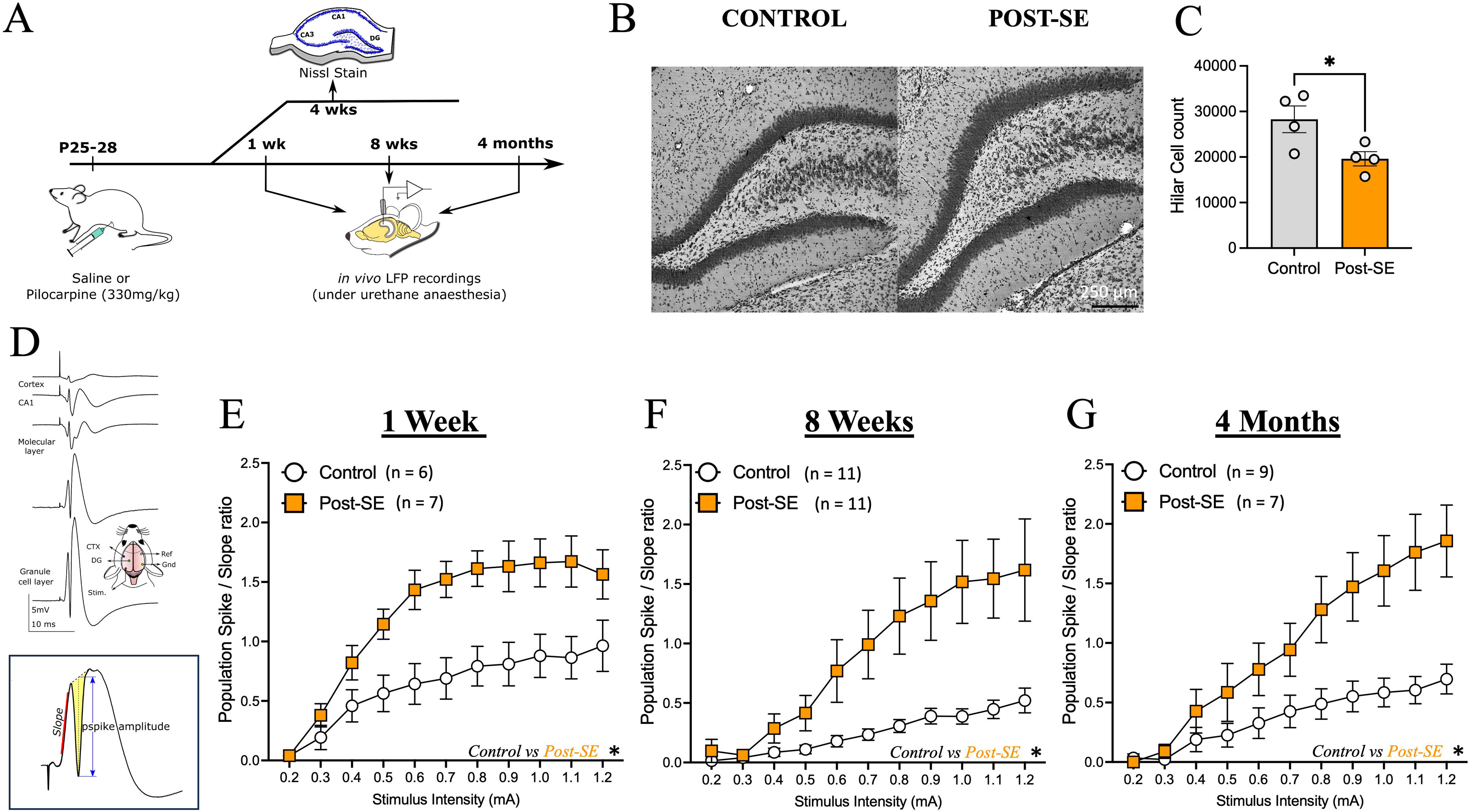
Pilocarpine-induced status epilepticus promotes hilar cell death and persistent network hyperexcitability. A. Schematic of experimental timeline. B-C. Representative Nissl stain images (B) and quantification (C) of hilar neurons from control and post-SE rats 4 weeks after pilocarpine-induced SE (N= 4 rats/group). D. Illustrations showing electrode depth/voltage profile of responses to perforant path stimulation. *Inset* shows the calculation of slope and population spike amplitudes. E-G. Summary population spike/ EPSP slope ratio, indicating an increase in network excitability at 1 week (E), 8 weeks (F) and 4 months (G) after SE compared to age-matched sham controls. * indicates *P*<0.05 by unpaired t-test (C) and TW RM ANOVA (E-G).

Excitation-spike (E-S) coupling, the probability that a given excitatory synaptic input drives an action potential, is known to be compromised or dysregulated in hippocampal CA1 during epileptogenesis (Carpenter-Hyland et al., 2017). However, the temporal progression of changes in DG E-S coupling following SE remains ill-defined. To quantify E-S coupling *in vivo*, we measured input-output relationships between the population spike amplitude and excitatory post-synaptic potential (EPSP) slope in urethane-anesthetized control and post-SE rats (Figure 1D, inset). Our experiments revealed a significant increase in population spike/ EPSP slope ratios as early as 1 week after SE, demonstrating an early enhancement of E-S coupling in the DG (N = 6 control, 7 post-SE rats; P = 0.0096 by two-way RM ANOVA, F (1,11) = 9.773; Figure 1E). This increase in E-S coupling persisted 8 weeks (N=11 control and 11 post-SE rats; *P* = 0.0066 by two-way RM ANOVA, F (1,20) =9.204, Figure 1F) and 4 months after SE (N=9 control and 7 post-SE rats, *P* = 0.0061 by two-way RM ANOVA, F (1,14) =10.41, Figure 1G). Together, these findings demonstrate an early, persistent, and long-term elevation of DG network excitability following SE.

To test whether altered inhibitory regulation within the DG contributes to the post-SE enhancement of E-S coupling, we evaluated feedback inhibition *in vivo* using paired-pulse stimulation across 20, 40, and 60 ms inter-pulse intervals (Figure 2A-B). Across all three inter-pulse intervals, two-way ANOVA revealed a robust main effect of group, indicating that SE significantly alters GABA_A_-mediated feedback inhibition (F (1,37) = 5.43, *P* =0.025 at 20 ms; F (1,37) = 26.56, *P* <0.0001 at 40 ms; and F (1,37) = 25.03, *P* <0.0001 at 60 ms; Figure 2C-E). At a 20 ms inter-pulse interval, when the paired pulse ratio (PPR) is heavily dominated by recurrent GABAergic feedback synaptic inhibition, PPR evolved dynamically across the post-SE timeline. Post-hoc comparisons revealed that the PPR at 20 ms inter-pulse intervals was comparable to controls at 1 week (*P* =0.8001) and 4 months (*P* =0.1053, Figure 2C) after SE, but was significantly decreased (reflecting enhanced feedback inhibition) at 8 weeks post-SE (*P* = 0.0418). In contrast, at 40 ms and 60 ms inter-pulse intervals, where control animals typically exhibit paired pulse facilitation, post-SE animals exhibited significant paired pulse depression at both 1 week (*P* =0.0002 and *P* <0.0001, respectively) and 8 weeks (*P* = 0.0026 and *P* = 0.0058, respectively; Figure 2D). At 4 months post-SE, we did not observe any difference in 40 ms or 60 ms PPR between compared matched controls 4 months (*P* = 0.1581 and *P* = 0.2324, respectively; Figure 2E).

**Figure 2:**
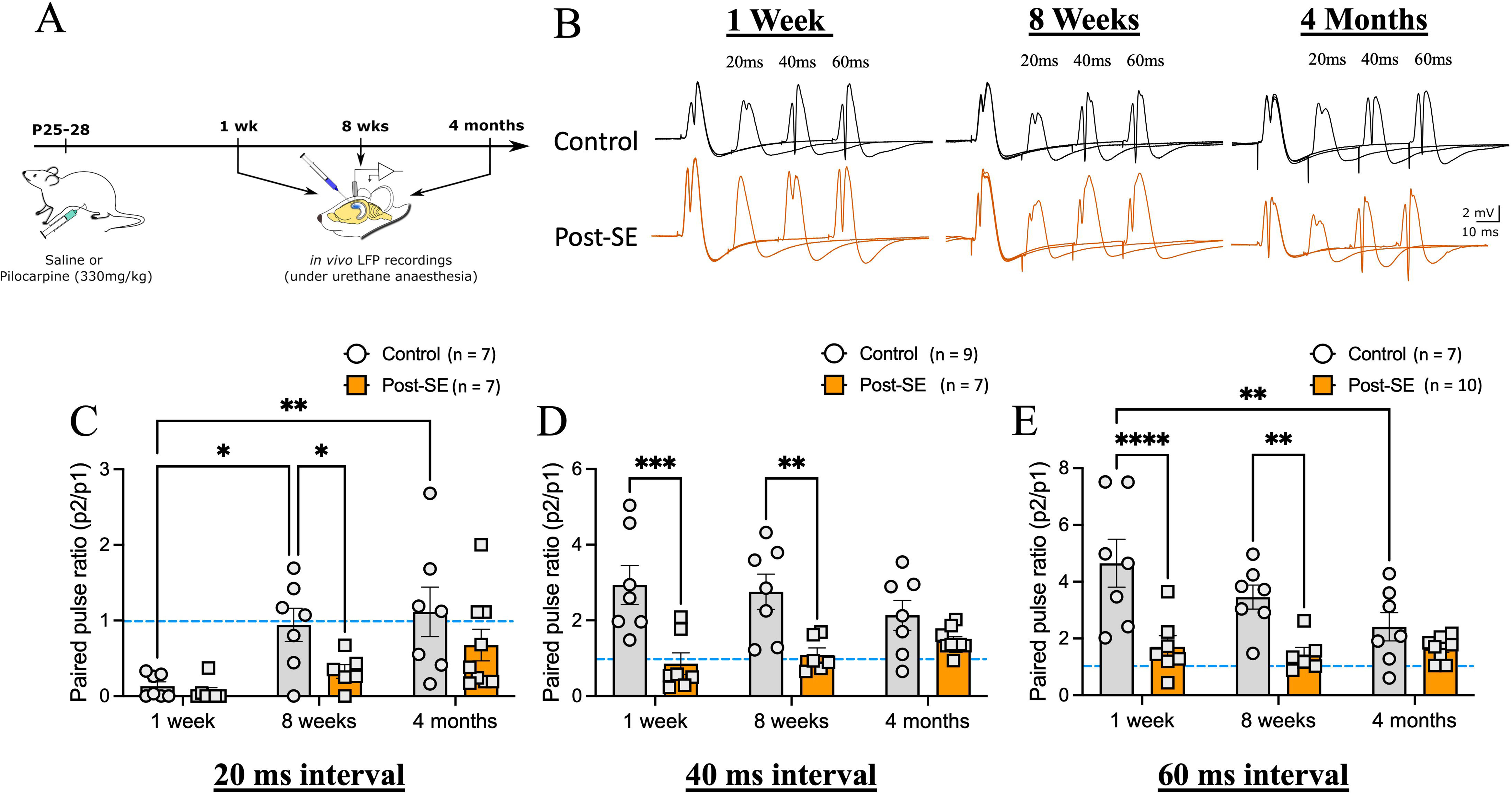
Time-dependent changes in dentate paired pulse ratio following pilocarpine-induced status epilepticus. A. Schematic of experimental timeline. B. Representative *in vivo* local field potential (LFP) traces evoked by perforant path stimulation in urethane-anesthetized control and post-SE rats show responses to paired electrical stimulations at 20ms, 40ms and 60ms inter-pulse intervals. C-E Summary of paired pulse ratios, recorded 1 week (B, C), 8 weeks (B, D) and 4 months (B, E) after SE induction. * indicates *P*<0.05 by TW RM ANOVA followed by Šídák’s multiple comparisons test.

Notably, epileptogenesis appeared to disrupt the normal age-dependent maturation of DG feedback circuits. In control animals, the 20 ms PPR exhibited a progressive age-dependent increase from 5 weeks (1 week post-saline) to 5 months (4 months post-saline, (F (2,37) =8.85, *P* = 0. 0007), reflecting a developmental modulation of feedback inhibition strength. This developmental trajectory was blunted in the post-SE animals (Figure 2C). Similarly, at the 60 ms inter-pulse intervals, control animals showed a significant decrease in PPR between 5 weeks and 5 months of age (*P* = 0.0052), a developmental transition that was entirely absent following SE. Notably, this developmental shift in control animals, rather than true physiological recovery in the post-SE group, accounted for the convergence of control and post-SE PPR values at 4 months. Finally, the lack of a significant group × age interaction across all inter-pulse intervals (20 ms: F (2,37) = 0.0954, *P* = 0.3944; 40 ms: F (2,37) =2.241, *P* = 0.1206; 60 ms: F (2,37) = 2.903, *P* = 0.0675 at 60 ms) indicates that while the magnitude of circuit disruption varies across disease progression, SE enhances dentate gyrus inhibitory control.

Because our findings demonstrated elevated E-S coupling alongside disruptions in circuit inhibition, we next investigated whether these alterations translate to enhanced network throughput *in vivo*. As a functional metric of DG throughput, we quantified changes in dentate spike (DS) frequency using 16-channel silicon probe recordings in urethane-anesthetized rats (Figure 3 A-B). Although early shifts in DS frequency have been reported after brain injury (Subramanian et al., 2026) and kainic acid-induced SE (Flynn et al., 2015), whether these changes persist long-term is currently unknown. Our recordings revealed a significant elevation in DS frequency 4 weeks after SE compared to age-matched controls (frequency in Hz, control: 0.11±0.01, post-SE: 0.25±0.04, N= 5 rats each, *P* = 0.0165 by unpaired t-test, t=3.022, df=8, Figure 3A-C). Crucially, our experiments revealed that this increase in DS frequency persisted chronically up to 10 months post-SE (frequency in Hz, control: 0.13±0.01, post-SE: 0.23±0.01, N= 6 control and 4 post-SE rats, *P*=0.0010 by unpaired t-test, t=5.040, df=8, Figure 2D). Together, these data demonstrate an early-onset, long-term elevation of DS frequency following pilocarpine-induced SE, pointing to sustained DG network hyperexcitability that may compromise downstream hippocampal signal processing and cognitive function.

**Figure 3:**
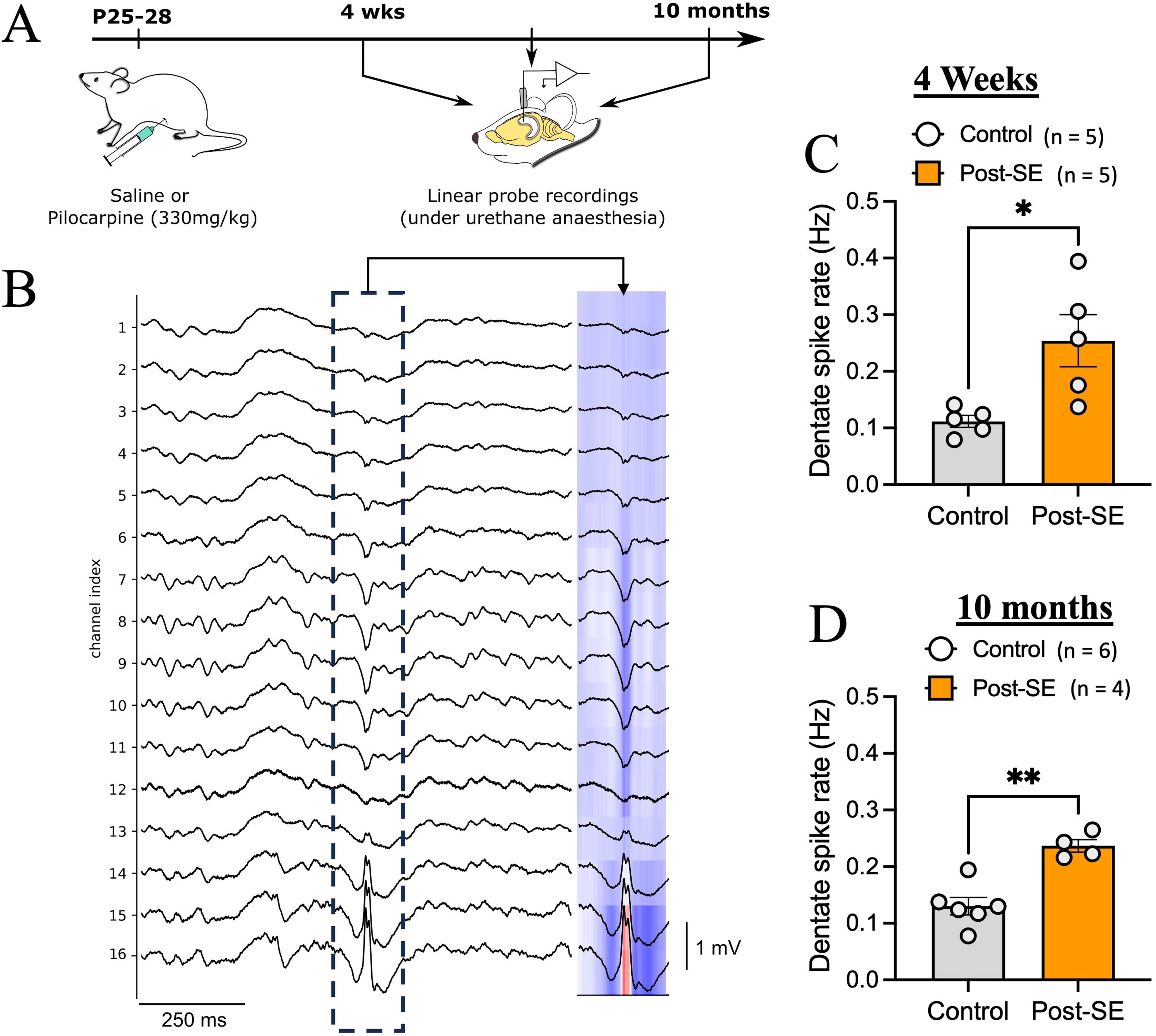
Persistent increase in dentate spike rates following pilocarpine-induced status epilepticus. A. Schematic of experimental timeline. B. Representative trace from 16 Ch linear probe recordings from urethane anesthetized rat highlighting dentate spikes. *Inset* shows current-source density plot of a dentate spike in the boxed area. C. Quantification of dentate spike frequency indicate a significant increase in frequency in rats 4 weeks after SE compared to age-matched controls. D. Quantification of dentate spikes from rats 10 months after SE induction. * indicates *P*<0.05, ** indicates *P*<0.01 by unpaired t-tests.

To evaluate whether post-SE alterations in E-S coupling, feedback inhibition, and network throughput are driven by presynaptic GABA_B_ receptor-mediated mechanisms, we sought to isolate presynaptic signaling within the DG circuit. Previous studies investigating presynaptic GABA_B_ receptor function have typically relied on non-selective antagonists that block both pre- and postsynaptic receptors (Mott et al., 1993; Mott and Lewis, 1991) or constitutive transgenic knockout models (Foster et al., 2013). However, systemic knockout strategies can introduce developmental or network-level compensatory confounds. To circumvent these limitations and directly test the contribution of presynaptic GABA_B_ receptor signaling *in vivo*, we used CGP36216, a selective presynaptic GABA_B_ receptor antagonist (Adkins et al., 2021; Li et al., 2024; Lynch et al., 2017; Ong et al., 2001; Yang et al., 2021).

To determine the contribution of presynaptic GABA_B_ receptors to DG E-S coupling, we performed simultaneous *in vivo* field recordings paired with local intrahippocampal injection of the selective presynaptic GABA_B_ receptor antagonist CGP36216 (3mM) using cannula electrodes (Figure 4A-B). While intrahippocampal CGP36216 failed to alter E-S coupling in control animals, it significantly enhanced E-S coupling in the post-SE group at 1 week (*P* = 0.009; N = 5 rats/group), and 8 weeks (*P* = 0.0171; N = 7 control and 6 post-SE rats, Figure 4C-E). By 4 months post-SE, CGP36216 no longer altered E-S coupling in either control or post-SE groups (Effect of Drug: F (1,8) = 3.744, *P* = 0.0890; N = 5 rats/group, Figure 4F). No significant drug × group interactions were observed across timepoints (p>0.05 for all). These findings demonstrate that early after SE, presynaptic GABA_B_ receptor signaling exerts a homeostatic braking effect that dampens aberrant E-S coupling, a compensatory mechanism that is no longer present 4 months post-SE (Figure 4A–F).

**Figure 4:**
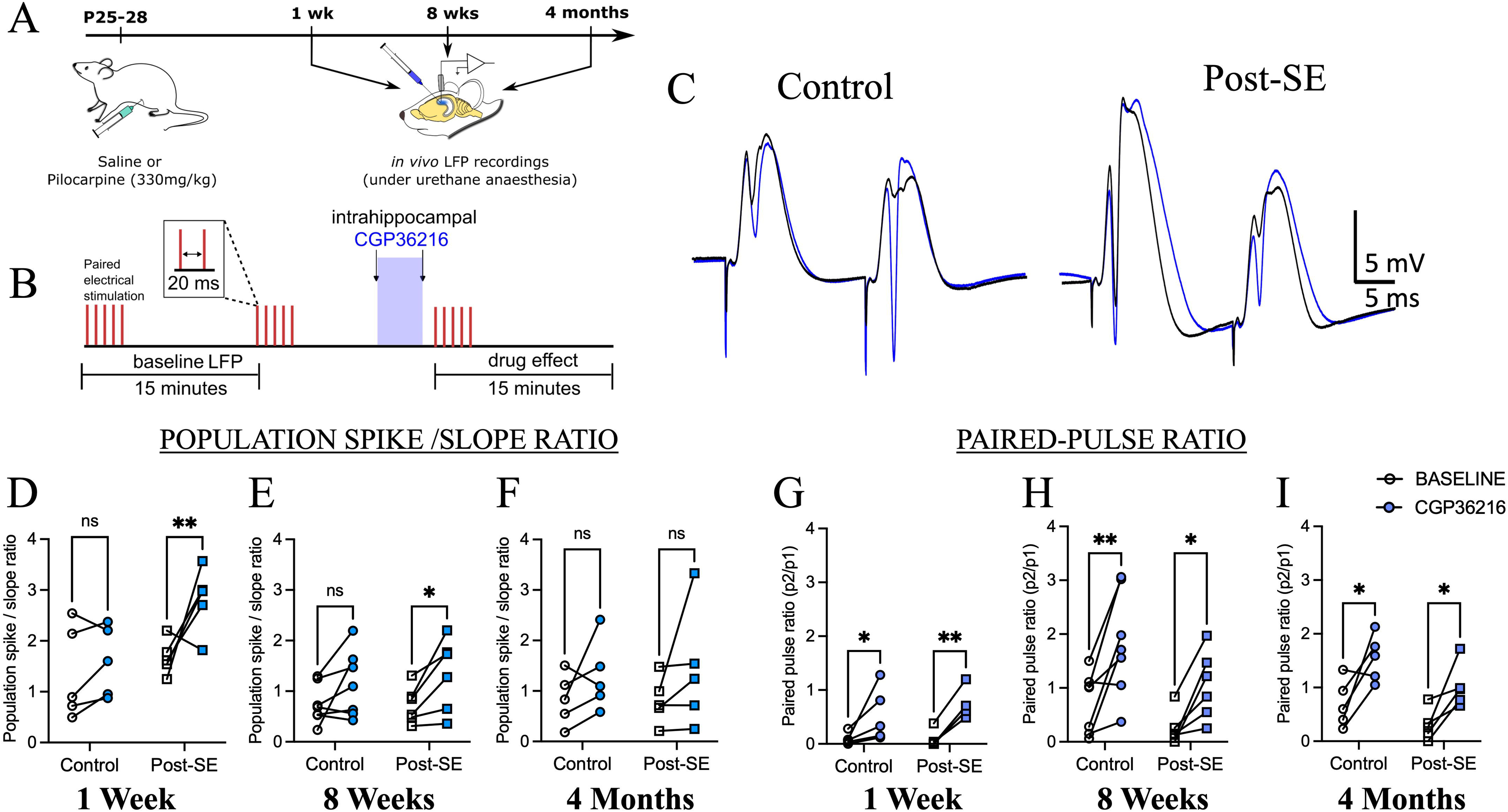
Selective presynaptic GABA_B_ receptor inhibition reduces network inhibition. A. Schematic of experimental timeline. B. Schematic showing how evoked and spontaneous LFPs were recorded before and after intrahippocampal infusion of CGP36216. C. Representative in vivo LFP traces show response to paired stimulation at 20ms. Recordings following intrahippocampal CGP36216 infusion are in blue. D-F. EPSP-spike coupling was enhanced by CGP36216 at 1 week (D), 8 weeks (E) after SE, but not at 4 months (F). G-I. CGP36216 increased paired pulse ratio at 20ms inter-pulse interval in both control and post-SE at all time points (G-I). * indicates *P*<0.05, ** indicates *P*<0.01 by TW RM ANOVA.

To evaluate the contribution of presynaptic GABA_B_ receptors to dentate gyrus feedback regulation, we next examined the effect of CGP36216 on PPR at 20 ms inter-pulse interval. Intrahippocampal CGP36216 perfusion significantly increased PPR across all post-SE and control time points, reflecting a reduction in feedback inhibition (Figure 4G–I). Two-way repeated-measures ANOVA showed a significant main effect of drug treatment at 1-week (F (1,8) = 25.9, *P* = 0.0009; N = 5 rats/group), 8-weeks (F (1,11)=23.47, *P* = 0.0005; N = 7 control and 6 post-SE rats), and 4 months (F (1,8)=18.0, *P*= 0.0028; N = 5 rats/group). No significant drug × group interactions were identified at any of the time points examined (p>0.05 for all). Mechanistically, these data indicate that presynaptic GABA_B_ signaling selectively limits auto-inhibitory feedback onto GABAergic interneurons, thereby sustaining recurrent inhibitory drive in both control and post-SE circuits. Combined with our E-S coupling findings, these results suggest that early after SE, presynaptic GABA_B_ receptors function to bolster inhibition and constrain aberrant E-S coupling.

Because hilar interneuron activity strongly correlates with DS generation (Penttonen et al., 1997), we next asked whether presynaptic GABA_B_ regulation of E-S coupling and feedback inhibition also influences overall dentate throughput. To achieve simultaneous electrophysiological recording and local drug injection, we utilized single-tungsten depth electrodes coupled to cannula probes (Figure 5A–B), rather than the multi-site silicon probes used in Figure 3. Comparing baseline DS rates recorded via tungsten electrodes versus multi-channel silicon probes revealed no technique-dependent differences in control basal DS frequency and reaffirmed the robust post-SE elevation in DS rates, validating our single-channel recording and detection pipeline (Frequency in Hz: 1-week: control 0.12± 0.02, post-SE-0.31±0.05, *P* = 0.0001, N = 5 control and 5 post-SE rats; 8-weeks: 0.09±0.01, post-SE-0.23±0.04, *P* = 0.0026, N = 11 control and 11 post-SE rats ;4-months: control-0.13±0.04, post-SE-0.38±0.06, *P*<0.0001, N = 8 control and 7 post-SE rats; Figure 5A-C). Furthermore, vehicle controls confirmed that the intrahippocampal injection itself did not disrupt network dynamics, as intrahippocampal saline infusions had no significant effect on DS frequency in either group (P>0.05 for all comparisons, Figure 5D).

**Figure 5:**
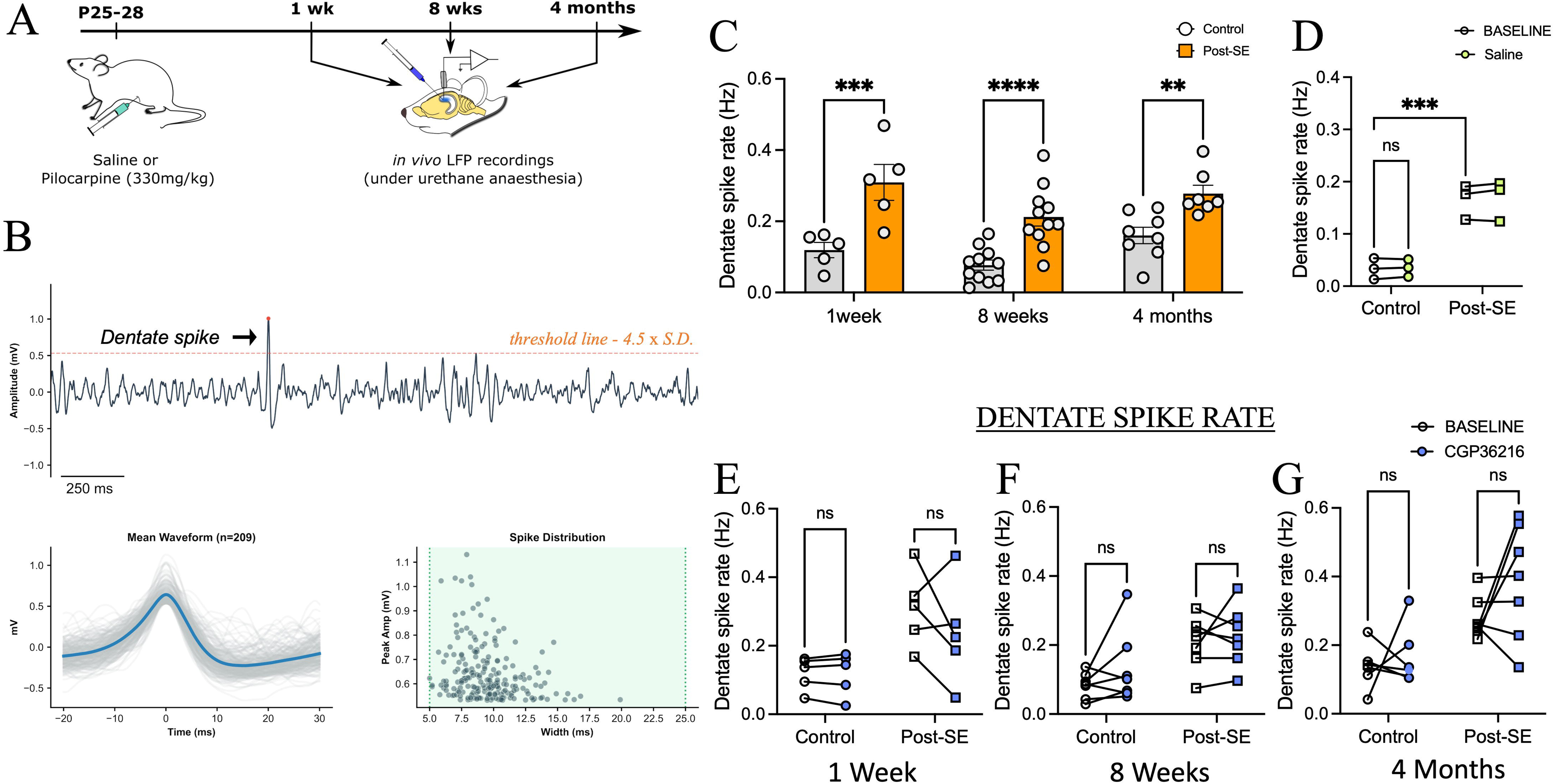
Inhibition of presynaptic GABA_B_ receptors does not alter dentate spike frequency. A. Schematic of experimental timeline. B. Representative in vivo LFP trace (top), average dentate spikes detected from an animal (lower-left panel,) and distribution of half-width and amplitude (lower-right panel). C. Quantification of dentate spikes recorded using single-depth electrode indicates an increase in dentate spike frequency at 1 week, 8 weeks, and 4 months post-SE. D. Intrahippocampal infusion of vehicle did not alter DS frequency. E-G. Intrahippocampal infusion of CGP36216 did not alter dentate spike frequency at all 3 time points. * indicates *P*<0.05, ** indicates *P*<0.01, *** indicates *P*<0.001, **** indicates *P*<0.0001 by TW ANOVA (C) or TW RM ANOVA (D-G).

In contrast to its prominent effects on E-S coupling and PPR, intrahippocampal CGP36216 infusion failed to alter DS frequency in any group or time point tested. Specifically, neither the main effects of drug or drug X group interaction reached significance at any of the three time points (1 week: (F(1,8)=1.323, *P* = 0.2833), N = 5 control rats and 5 post-SE rats; 8-weeks - F (1,12) = 2.126, *P* = 0.091, N= 7 control and 7 post-SE rats; 4 months- (F(1,11)=2.213, *P*=0.1649, N=5 control and 7 post-SE rats; Figure 6E-G). These data indicate that while presynaptic GABA_B_ receptors dynamically regulate monosynaptic E-S coupling and local feedback inhibition, they do not directly gate the network mechanisms driving spontaneous DS frequency.

**Figure 6:**
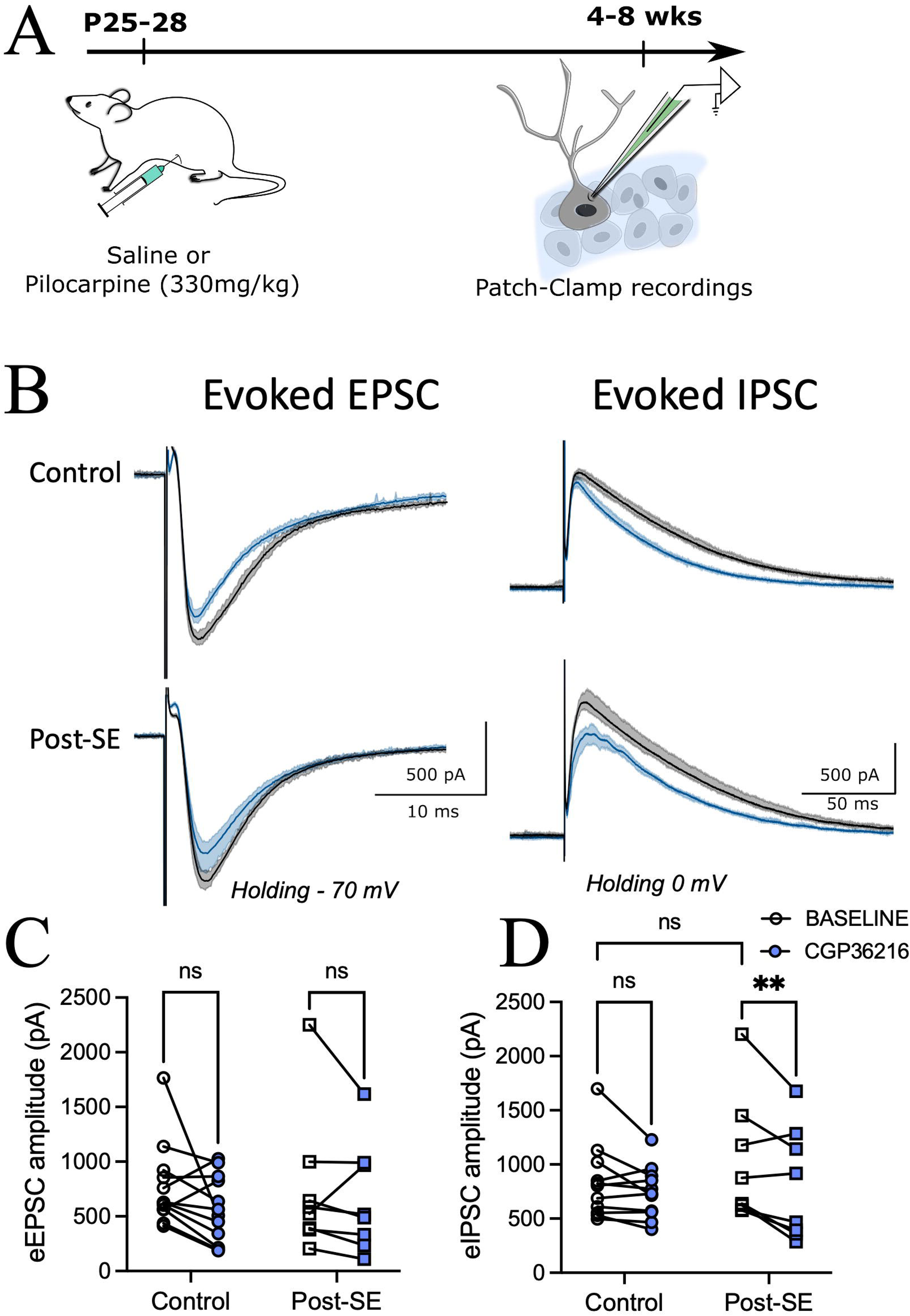
Inhibition of presynaptic GABA_B_ receptors selectively reduces IPSC amplitude in dentate granule cells after SE. A. Schematic of experimental timeline. B. Representative perforant path-evoked EPSC traces recorded in slices from control (above) and post-SE rats (below)from a holding potential of -70 mV (left panels). Responses during perfusion CGP36216 (100µM), are overlaid in blue. Example perforant path-evoked IPSC traces recorded in slices from control (above) and post-SE rats (below) from a holding potential of 0 mV (right panels). Responses to CGP36216 (100µM) are overlaid in blue. C-D. Quantification of the amplitude of evoked EPSC (C) and evoked IPSC (D) before and in the presence of CGP36216 (blue). ** indicates *P*<0.01 by paired t-test (D), TW RM ANOVA.

Finally, to establish a cellular mechanism for the unexpected CGP36216-mediated increase in PPR, we evaluated presynaptic GABA_B_ receptor regulation of excitatory and inhibitory synaptic inputs to dentate granule cells using whole-cell patch-clamp recordings in acute hippocampal slices from rats 1 week after saline or pilocarpine-SE. Bath perfusion of CGP36216 (100µM) did not alter evoked glutamatergic EPSC (eEPSC) amplitude in either control or post-SE animals (N=12 cells from 5 control rats, 8 cells from 4 post-SE rats, Effect of drug *P* = 0.0840 by two-way RM ANOVA F (1,18) =3.346, Figure 6A-C). In contrast, CGP36216 significantly reduced evoked IPSC (eIPSC) amplitude in DGCs at 1 week post-SE (N=11 cells from 5 control rats, 8 cells from 4 post-SE rats, Effect of drug P = 0.0037 by Two-way RM ANOVA F (1,17) =11.26, Figure 4D). Because a presynaptic GABA_B_ receptor antagonist decreased, rather than increased, eIPSC amplitudes in DGCs after SE, these data suggest a polysynaptic disinhibitory role for presynaptic GABA_B_ receptors after SE. Specifically, the data suggest that endogenous activation of presynaptic GABA_B_ receptors after SE suppresses interneuron-to-interneuron inhibitory transmission, disinhibiting feedback interneurons and thereby bolstering feedback inhibition onto DGCs. These cellular findings directly align with prior evidence demonstrating a selective post-SE increase in presynaptic GABA_B_ signaling at synapses connecting cannabinoid receptor type-1 (CB1)-expressing interneurons to parvalbumin-positive basket cells (Yu et al., 2016), providing a mechanistic basis for the observed changes in PPR and network E-S coupling.

## DISCUSSION

Structural and functional reorganization of the DG network is a classic hallmark of temporal lobe epilepsy (Dengler and Coulter, 2016; Dudek and Sutula, 2007; Mello et al., 1993; Scharfman, 2019; Sloviter, 1994a). While the failure of DG to effectively filter afferent inputs is believed to drive hippocampal hyperexcitability and seizures, how the DG network dynamics evolve over time after SE is still not fully understood. Our findings demonstrate a significant and persistent increase in E-S coupling, feedback inhibition, and dentate spike frequency, indicating long-lasting alterations in DG function after SE. Importantly, our experiments support a transient role for presynaptic GABA_B_ receptors in constraining the emergent hyperexcitability in the DG network after SE.

Distinct populations of hilar inhibitory neurons are vulnerable to SE, and have been associated with reduced inhibitory control of DG granule cells (Kobayashi and Buckmaster, 2003; Obenaus et al., 1993; Sloviter, 1987). In line with this, we found a significant reduction in hilar cell population in rats post-SE, which correlates with the observed increase in network excitability (Figure 1). We find that *in vivo* excitation-spike coupling, an integrated measure of network excitability (Abraham and Bliss, 1985; Carpenter-Hyland et al., 2017) was significantly increased post-SE across all timepoints examined, indicating post-SE changes in granule cell excitability and feedforward inhibition. Our data demonstrating lack of CGP36216 modulation of E-S coupling in controls is consistent with studies using GABA_B_ agonists and antagonists that modulate both pre- and post-synaptic receptors, which show limited basal GABA_B_ regulation of single-stimulus-evoked responses (Otis et al., 1993; Otis and Mody, 1992). However, the ability of CGP36216 to enhance E-S coupling for up to 8 weeks after SE indicates the presence of a basal presynaptic GABA_B_ tone that restrains DG network excitation after SE.

While several studies have examined the temporal dynamics of changes in fast GABA_A_ receptor-mediated synaptic inhibition in experimental epilepsy, few studies have examined how alterations in GABA_B_ receptor signaling affect dentate circuit function after SE (Avoli and Lévesque, 2022). GABA_B_ receptors have both pre- and postsynaptic functions; postsynaptic GABA_B_ signaling leads to slow inhibitory potassium currents, while presynaptic GABA_B_ signaling can mediate homo- and heterosynaptic suppression of synaptic release at GABAergic and glutamatergic terminals (Gassmann and Bettler, 2012). Current knowledge on GABA_B_ receptor function relies heavily on nonselective pharmacology that cannot selectively modulate pre- versus postsynaptic receptors. The recent availability of a GABA_B_ antagonist selective for presynaptic receptors (Adkins et al., 2021; Li et al., 2024; Lynch et al., 2017; Ong et al., 2001; Yang et al., 2021) enabled us to evaluate the role of presynaptic GABA_B_ signaling in regulation of basal dentate excitability and feedback inhibition both in controls and over the time course of disease progression after SE.

Based on our prior study identifying a selective increase in presynaptic GABA_B_ signaling in CB1-positive interneuron synapses onto PV-positive basket cells after SE (Yu et al., 2016), we propose that presynaptic GABA_B_ receptors play a disinhibitory role after SE by suppressing inhibitory synapses to perisomatic PV-positive basket cells, thereby increasing inhibition of granule cells (Foster et al., 2013; Yu et al., 2016) (Figure 7, left panel). This proposal is also consistent with literature showing selective increase in GABA_B_ receptor expression in dentate hilar inhibitory neuron subtypes after SE (Foster et al., 2013; Straessle et al., 2003). Since PV-basket cells are activated more rapidly than granule cells in response to perforant path stimulation, they are well positioned to regulate E-S coupling through feedforward inhibition (Bartos and Elgueta, 2012; Pelkey et al., 2017). The increase in basal PV-mediated inhibition is a potential compensatory mechanism to restrain the increase in E-S coupling after SE (Hansen et al., 2018). Our data showing that blocking presynaptic GABA_B_ signaling reduces evoked IPSC amplitude in post-SE rats without impacting evoked IPSC amplitude controls is consistent with the proposed disinhibitory mechanism and a basal increase in presynaptic GABA_B_ tone after SE (Figure 7, right panel). Moreover, CGP36216 failed to alter evoked EPSC amplitude in granule cells from both controls and post-SE rats, demonstrating that GABA_B_ receptors do not modulate heterosynaptic glutamatergic release at perforant path terminals.

**Figure 7:**
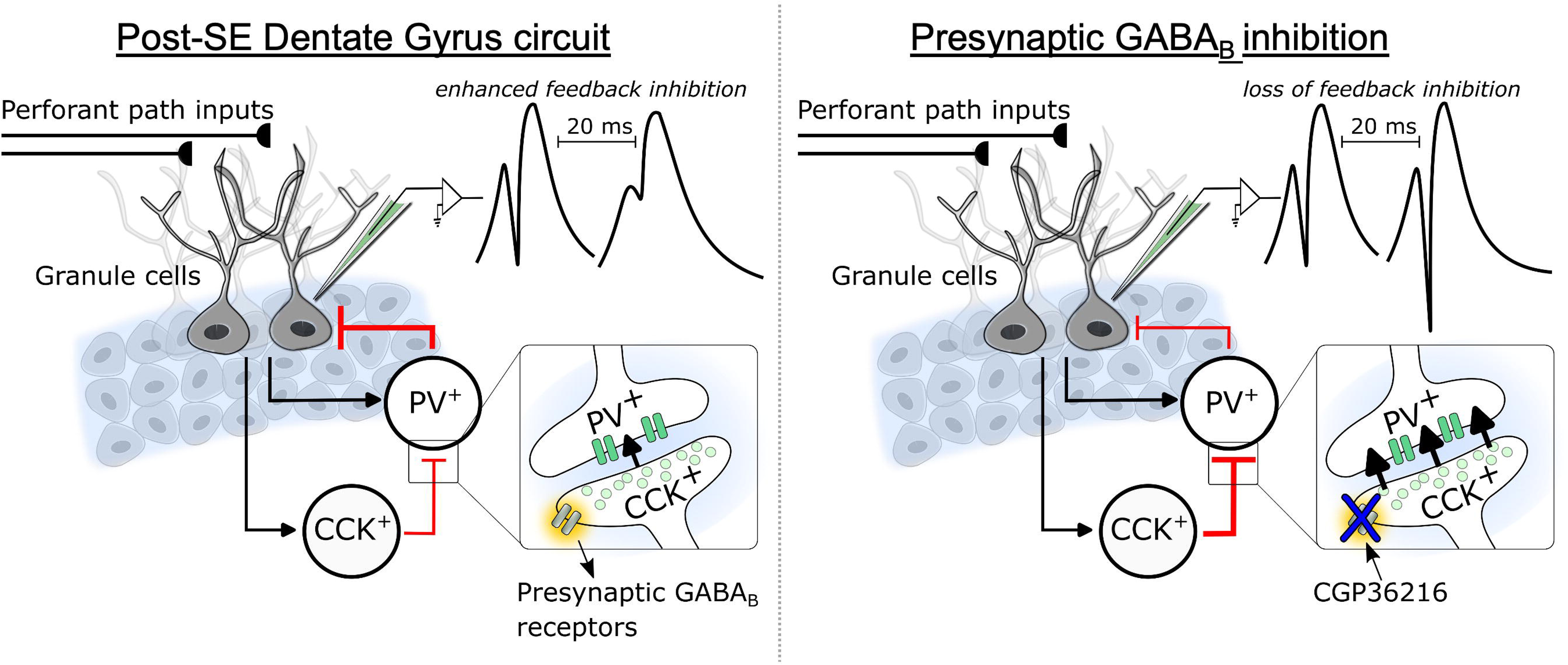
Proposed mechanism for disinhibition of DG granule cells by presynaptic GABA_B_ receptors after SE. Summary figure of proposed mechanism that leads to enhanced synaptic inhibition after pilocarpine induced SE. Following SE, the enhanced presynaptic GABA_B_ tone reduces synaptic release from CCK+ interneuron to PV+ interneurons (Based on findings in Yu et al 2015). This disinhibition of PV neurons enhances perisomatic inhibition on dentate granule cells (left panel). Inhibition of presynaptic GABA_B_ receptors (in current study) is hypothesized to maintain CCK interneuron to PV interneuron inhibition and reduce PV+ interneurons mediated feedback inhibition of granule cells.

Paired pulse depression at short interstimulus intervals is typically attributed to fast feedback inhibition by PV-positive interneurons (Jedlicka et al., 2010). As with prior studies in kindling models of epileptogenesis (De Jonge and Racine, 1987; Stringer and Lothman, 1989) and human epileptic tissue (Masukawa et al., 1996; Wilson et al., 1998), we find an increase in paired pulse inhibition up to 8 weeks post-SE, indicating enhanced feedback inhibition. It is interesting that the paired pulse ratios at 20 ms and 60 ms show significant age-related changes within the control rats (Figure 2C-E), which are not seen after SE. CGP36216 consistently and robustly increased PPR measured at 20 ms intervals in all groups, indicating a key role for presynaptic GABAB receptors in regulating feedback inhibition. These findings suggest that presynaptic GABA_B_ activation can act at the timescale of recurrent feedback inhibition and enhance inhibition. Taken together, it is worth speculating that the GABA_B_- dependent disinhibitory circuit mechanism proposed above could also regulate polysynaptic feedback inhibition.

The chronic increase in dentate spike (DS) frequency is one of our study’s most notable findings. Recent publications have shown an early increase in DS frequency following kainic acid induced SE (Flynn et al., 2015) and brain injury (Subramanian et al., 2026). The present study provides the first evidence that DS frequency remains elevated into the chronic phase of epilepsy. The persistent increase in DS frequency may reflect the combined effects of altered entorhinal input, intrinsic granule-cell excitability, recurrent mossy-fiber remodeling, hilar cell loss, and changes in interneuron recruitment (Bragin et al., 1995; Penttonen et al., 1997). The chronic persistence of this phenotype despite normalization of baseline feedback inhibition at 4 months indicates that DS generation becomes increasingly dependent on distributed network properties that are only partially controlled by the inhibitory mechanism identified in the present study. The absence of an effect of CGP36216 on DS frequency is particularly informative, as it shows that E-S coupling does not determine DS frequency. This dissociation emphasizes the distinction between local input–output transformations and coordinated network events. While the mechanism of DS increase after SE remains to be determined, the persistent increase in DS identified here could contribute to the chronic cognitive comorbidities associated with epilepsy (Ewell, 2017; Farrell et al., 2024; Lensu et al., 2019; Nokia et al., 2017).

Collectively, our observations have implications for understanding therapeutic windows during epileptogenesis. Pharmacological enhancement of GABA_B_ signaling has been shown to suppress abnormal hippocampal activity, whereas GABA_B_ receptor antagonism can increase pathological discharges in experimental epilepsy (Avoli and Lévesque, 2022; Mareš et al., 2013; Rogawski and Löscher, 2004; Schuler et al., 2001; Smirnova et al., 2020). Our data suggest that presynaptic GABA_B_ receptors may represent a modulatory target during the early period when inhibitory circuitry retains substantial compensatory capacity.

